# Unprecedented Microviridae phages isolated from equine uteri

**DOI:** 10.1101/2025.03.06.640271

**Authors:** Lulu Guo, G. Reed Holyoak, Udaya DeSilva

## Abstract

Descriptions of the virome has only recently returned to the fore. There is much information yet to be discovered. Single-stranded DNA (ssDNA) viruses as an exceptionally widespread virus can infect diverse hosts. The simple genome of ssDNA viruses leads to a high propensity for mutation and recombination, which often pose infectious threats to society. However, research investigating ssDNA phage genomes have ignored the reproductive tract communities. Here we have identified novel members of the Microviridae family from horse (Equus cabalas) uteri, with completed genomic sequences. Comparative and phylogenetic analyses between different animals indicate that genetically similar animals to the equine likely have the same or similar viruses in their uterine microbiome. This research not only expands the understanding of the diversity within Microviridae family, but also provides new insights into potential therapeutics for uterine infectious disease.

## Introduction

When we first recognized the existence of microbes, we did not know that these tiny organisms are the cornerstone of the earth’s ecosystem and a vital factor in our health. Taking advantages of these commensal symbionts would greatly help us overcome the challenges we face today. As the mystery of the human genome has been gradually unveiled, the microbiome has become the focus of life science research and competitive marketing among countries. With the development of high-throughput DNA sequencing, microbial investigation has evolved from the role of a single pathogen in disease using Koch’s postulates to the study of an entire integrated microbial community. Diverse bacteria, archaea and microbial eukaryotes in the composition of a microbiome, including the pathobionts and pathogenic organisms, affect the health of the metaorganism through nutrition and drug metabolism (Wallace et al., 2010; de Clercq et al., 2016), synthesis of essential vitamins (Kau et al., 2011), defense against pathogens (Abt and Pamer, 2014), bile acid secondary processing (Wahlström et al., 2016), immune regulation (Round and Mazmanian, 2009; Belkaid and Hand, 2014), resistance to infection and susceptibility (Buffie et al., 2015), and even behavioral changes (Dinan et al., 2015). Nonetheless, it is now realized that viruses also play essential roles within a holobiont, both directly by infecting animal host cells and indirectly by modulating microbiome dynamics (Creasy et al., 2018).

As the life form with the most extensive distribution, the largest biomass, and the richest biodiversity on earth, viruses contain extremely rich species and genetic resources affecting the biosphere. A comprehensive and systematic analysis of viruses and a clear understanding of the relevant regulatory mechanisms will bring us innovative ideas and solutions in specific disease conditions. Viruses can infect not only human, animals, plants, but also bacteria. Bacteriophages are a type of virus that specifically infects bacteria. Phagein, a Greek word meaning to eat or devour, was the reference for coining the term bacteriophage by a French microbiologist named Félix d’Hérelle on September 3^rd^ in 1917. After 2 years, he applied phage therapy on a boy who was suffering from severe dysentery, and he became the first patient healed by using phage therapy. During the following years, doctors and researchers developed a passion for phage therapy, and successfully tested it against a variety of infectious diseases. Furthermore, Dr. D’Herelle suggested using a “phage cocktail” containing different phage strains, since bacteria develop resistance to a single phage. It is said that all medications have side effects. However, it appears that bacteriophage do not. Unlike antibiotics that will kill both pathogens and beneficial commensal bacteria, phages have affinity for specific receptors. Moreover, in contrast to the fact that the systemic concentration of antibiotics will decrease over time, each phage can produce ∼100 new phages through its replicative process (Campbell, 2003), which exponentially supports the attack of the host bacteria. They not only look like a robot, but also have their own “firewall-like” protection system. They will not start their “lytic programming” if they mis-attach to a bacterium rather than the target, and since they only survive within the host bacteria, they will die with the elimination of all the target pathogens.

The goal for viral metagenomics is to discover the viral diversity in the environment, which is often missed in studies targeting specific potential animal and bacterial hosts. Although both single-stranded DNA (ssDNA) and double-stranded DNA (dsDNA) phages have been found in animals (Wales et al., 2015; Malathi and Renuka Devi, 2019), the diversity of dsDNA phages is still the best characterized due to their large representation in cultures and genome sequence databases (Creasy et al., 2018). Also, commonly used library construction kits and virome protocols using linker adapter-amplified shotgun sequencing are not biased towards ssDNA viruses (Roux et al., 2016).

ssDNA viruses can be divided into 2 bacteriophage families, Inoviridae and Microviridae (King et al., 2011), and 9 families of eukaryotic viruse. Recent publications have shown that Microviridae seem to be ubiquitous. Microviridae were detected in marine environments (Mizuno et al., 2016), freshwater habitats (Roux et al., 2012a), ciona robusta (Creasy et al., 2018), stromatolites (Desnues et al., 2008), human gut environment (Roux et al., 2012b), sewage and sediments (Hopkins et al., 2014), dragonflies (Rosario et al., 2012), soil (Quaiser et al., 2015b; Williamson et al., 2017) and even integrated into the genomes of bacteroidetes species as temperate phages (Krupovic and Forterre, 2011). These studies markedly contributed to Microviridae specific genomic information, and enable more precise comparative Genome Analysis (Quaiser et al., 2015a).

The main purposes of this study were to: 1. obtain the full sequence of the novel Microviridae family from the endometrial lavage samples; 2. evaluate if they are common inhabitants of equine uterus; 3. construct phylogenetic tree and predict the probable host and lifestyle for them; 4. determine if they are unique to the horse. In this research, we report two novel ssDNA viruses from equine uteri. The new viral genomes presented in the study provide insights of the uterine environment and contribute to the documentation of Microviridae diversity.

## Materials and Methods

### Sample Collections and Processing

Equine uterine viral DNA was extracted from uterine lavage samples following a careful procedure as reported previously by our group (Reference https://rdcu.be/cZkV4). Using a similar protocol, bovine uterine lavage samples were from Oklahoma State University CVM Ranch cattle. Feline and canine uteri were aseptically collected from surgically obtained ovariohysterectomy specimens from the Oklahoma State University Boren Veterinary Medical Teaching Hospital. Porcine and ovine uteri were obtained avoiding contamination from carefully eviscerated animals from the Oklahoma State University Robert M. Kerr Food and Agricultural Products Center. Endometrial lavage samples were obtained aseptically first by ligating the excised end of the uterus. A sterile catheter was aseptically inserted into the uterine lumen and sterile saline was infused until the lumen was dilated. The infusate was then aspirated and stored at -80C until analyzed (Fig. 1). DNA was then extracted using Qiagen DNA Mini Kit (Qiagen, Inc., Valencia, CA, USA). NanoDrop 1000 Spectrophotometer (NanoDrop Technologies, Wilmington, DE, USA) was used to detect the DNA concentration.

**Figure 1:**
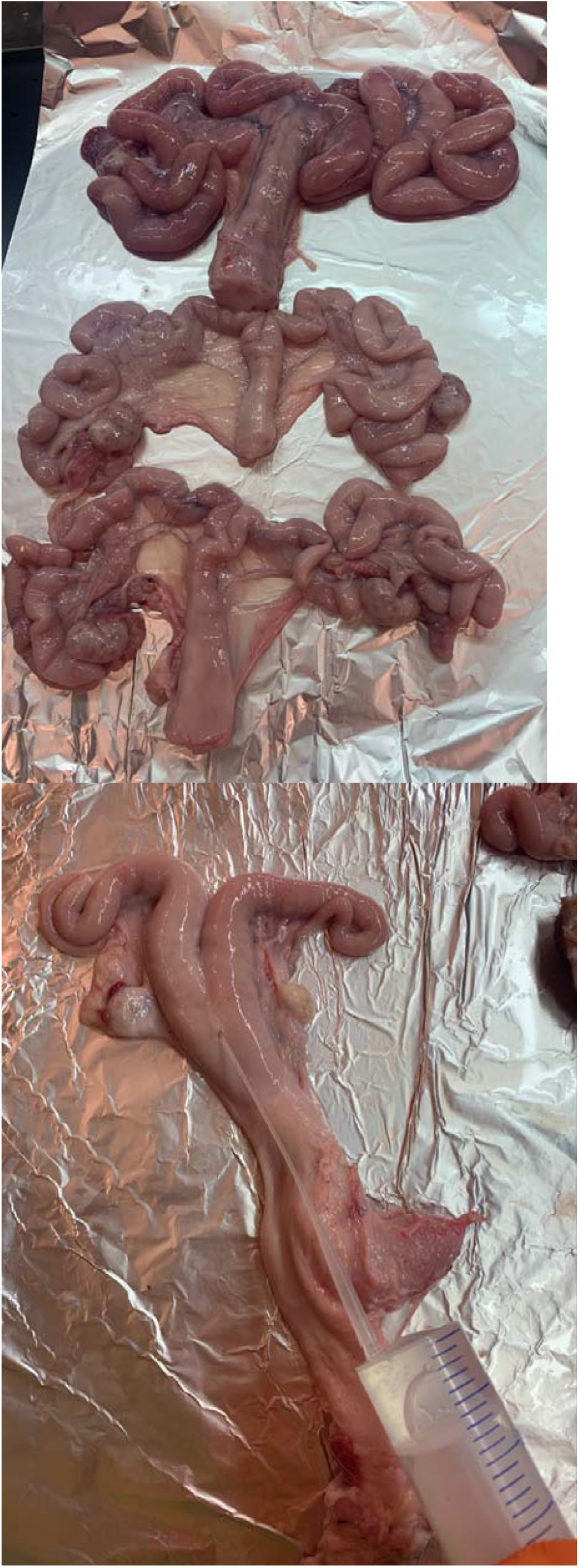
Biopsy samples treatment. Uterine lumens are flushed by saline with sterile catheter.

### Complete Contigs

Circular contigs sequence were determined by VIBRANT (Kieft et al., 2020). Blastx was then used to find Microviridae major capsid protein amino acid sequence in the circular contigs sequence. To avoid false positives, Blastn filtered the contigs to specified cellular organisms. In order to obtain the complete sequence of the novel Microviridae contigs, primers were designed and ordered accordingly by using Primer3 (Koressaar et al., 2018) and Integrated DNA Technologies, Inc. from both 3’ end and 5’ end. DNA was amplified in a 25μl PCR using 2.5μl 10X Buffer (Invitrogen, Waltham, MA, USA), 1.5μl 25μM Magnesium chloride (Invitrogen, Waltham, MA, USA), 0.5μl 10μM dNTP (Invitrogen, Waltham, MA, USA), 0.25μl 5U/μl Taq (Invitrogen, Waltham, MA, USA), 1μl template DNA, 1μl 10 μM forward primer (Integrated DNA Technologies, Inc. Coralville, IA, USA), 1μl 10 μM reverse primer (Integrated DNA Technologies, Inc. Coralville, IA, USA) and 17.25μl molecular grade water (Mediatech, Inc. Manassas, VA, USA) under the following conditions: 95°C for 1 minutes, followed by 35 cycles of 92°C for 30 seconds, 53°C for 30 seconds, and 72°C for 20 seconds. Afterwards, a final elongation step of 72°C for 2 minutes was performed. A 1.5% agarose gel was used to determine the success of PCR amplification. DNA was purified using PureLink™ Quick Gel Extraction Kit (Thermo Fisher Scientific Inc. Waltham, MA, USA). The purified DNA were sent to Oklahoma State University Center for Genomics and Proteomics for sanger sequencing. Results and the original contigs sequence were aligned with Clustal Omega (Sievers and Higgins, 2021) to make up the missing sequence.

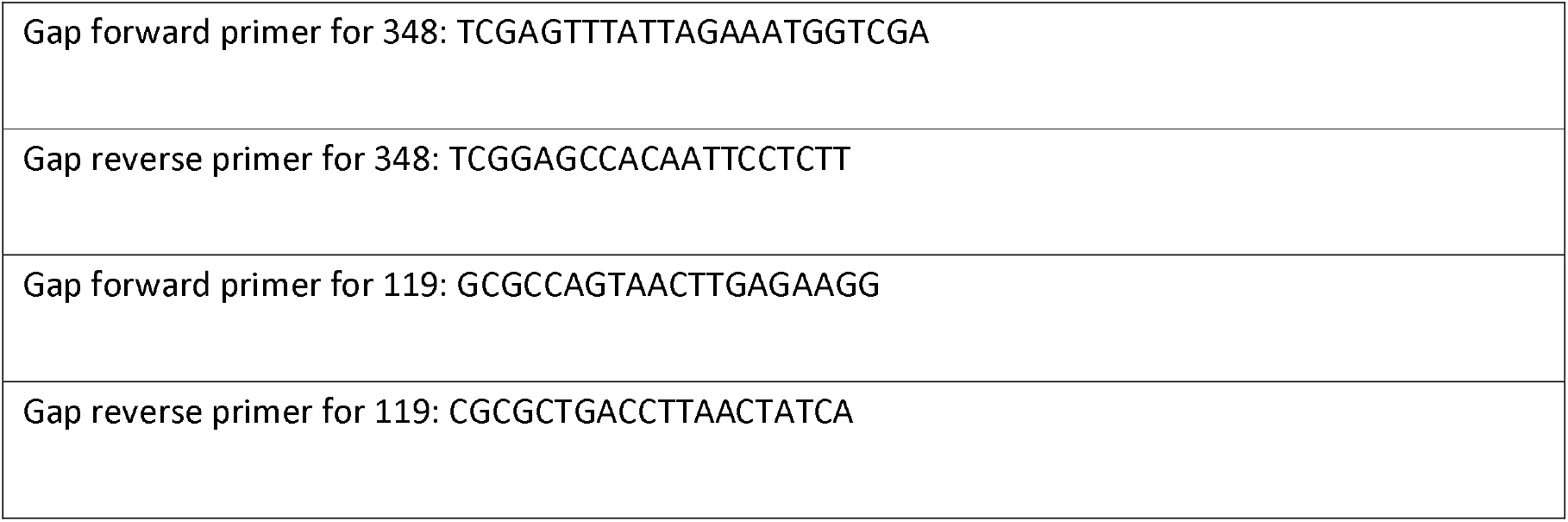

### Microviridae Genome Bioinformatic Analysis

Complete Microviridae genome were annotated using Prokka (Seemann, 2014). CGview (Stothard and Wishart, 2005) were then performed to get a circular map. Reference Microviridae family sequences were collected from NCBI. Phylogenetic trees were then created and visualized by Matlab. To predict the bacterial host, Prokaryotic virus Host Predictor (Lu et al., 2021), which is based on the differences of k-mer frequencies between viral and host genomic sequences was utilized. With a deep learning software called DeePhage (Wu et al., 2021), we analyzed the lifestyle of the novel Microviridae genome.

### Quantitative PCR Comparison

Primers were designed by Primer3 (Koressaar et al., 2018) using the equine viral assembly contigs 348 and 119. They were tested in equine samples first before moving forward to other animals to ensure the sanger sequencing results were the same as the assemble sequence. Primers were ordered from Integrated DNA Technologies, Inc. from both 3’ end and 5’ end. Five DNA samples of ovine, porcine and bovine, respectively, ten DNA samples of feline and canine, respectively, from the previous step were amplified in a 20μl qPCR reaction including 0.2μM Forward Primer, 0.2μM Reverse Primer, 10μl 2X supermix (Bio-Rad Laboratories, Inc. Hercules, CA, USA), 2μl DNA product and 6μl molecular grade water (Mediatech, Inc. Manassas, VA, USA). qPCR assays were performed with the thermo cycling profile comprising 1min initial denaturation at 98°C followed by 40 cycles of

95°C for 20s, 51°C/52°C, for 30s, melt curve analysis with CFX Connect Real-Time PCR Detection System (Bio-Rad Laboratories, Inc. Hercules, CA, USA) default setting. Three replicates of each sample genome including negative controls and positive controls (equine sample) were run together to identify mean and standard deviation of copy numbers. For the samples that showed positive in qPCR, ExoSAP-IT™ PCR Product Cleanup Reagent (Thermo Fisher Scientific Inc. Waltham, MA, USA) was used to remove the extra primers and qPCR reagents. Then the clean DNA were sent to Oklahoma State University Center for Genomics and Proteomics for sanger sequencing to check the correctness of sequence.

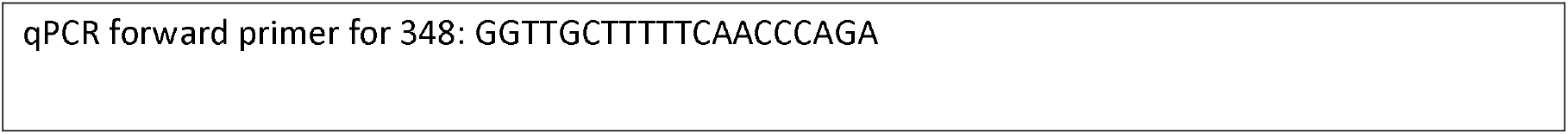
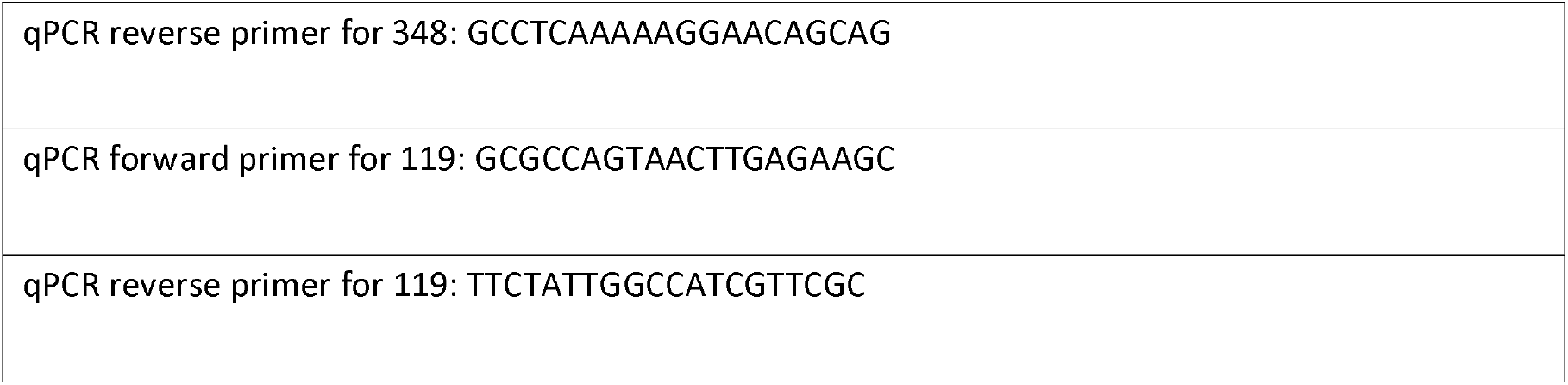

## Results

### Analysis of Complete Microviridae-Like Genomes

Contigs corresponding to complete circular genomes and high sequence similarity to the major capsid protein VP1 of Microviridae were obtained from the previous research by our team. In total, 2 new complete bacteriophage genomes were retained. To assess the completeness of the new Microviridae-affiliated genomes, PCR and sanger sequencing were performed using the forward primer from the end of the sequence and reverse primer from the beginning of the sequence. All primers bound at the expected position. Gap were filled by aligning sequences (Supplementary 1). Circular genome map (Fig. 2) shows the annotations of the 2 new viral genomes.

**Figure 2:**
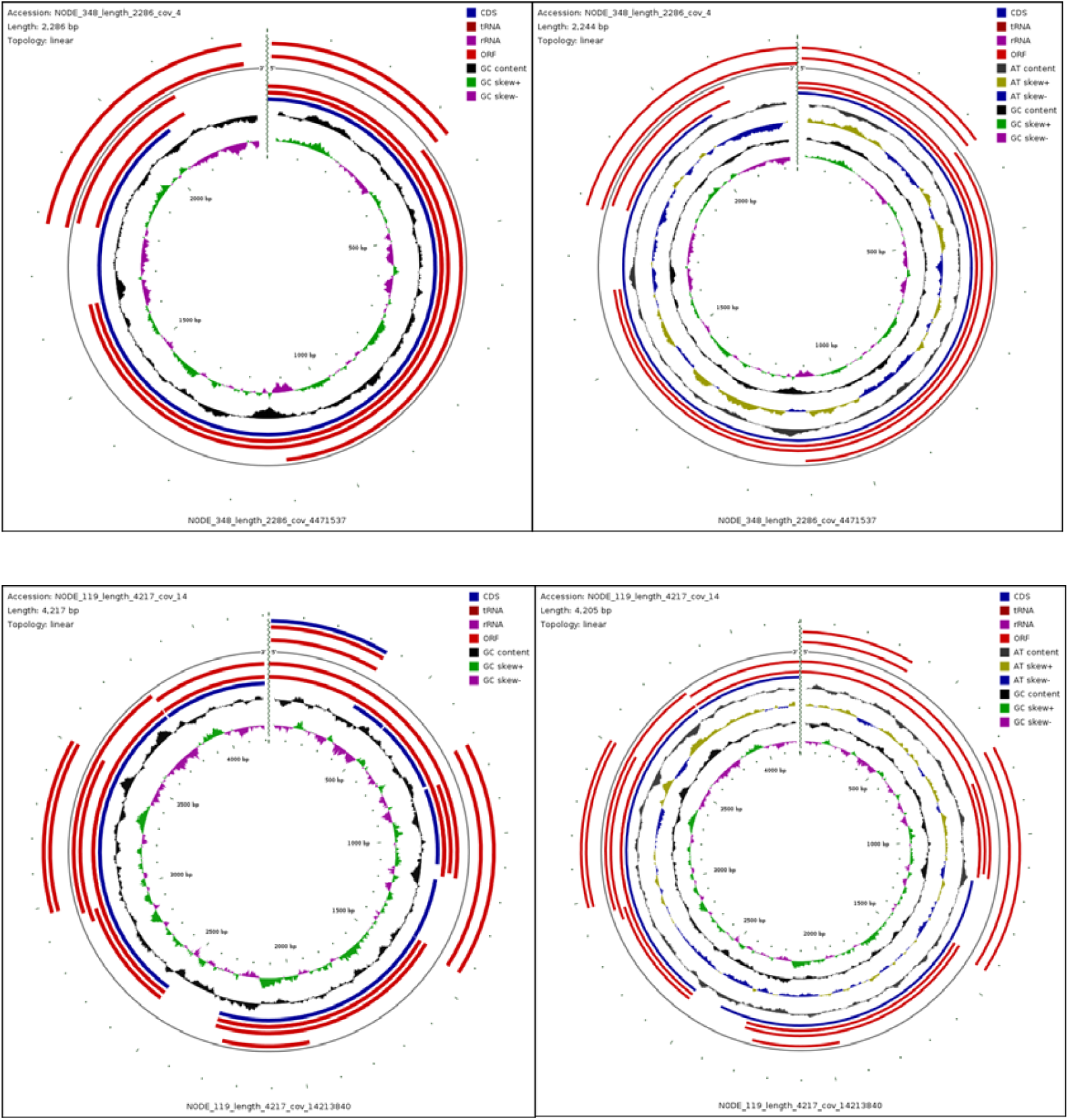
Circular genome map. Left side are maps before completing the sequence (original contigs) and after replenishing sequence with sanger sequencing results.

### Phylogenetic Analysis of the Conserved Major Capsid Protein VP1 with Known Microviridae Family

Phylogenetic analysis used the major capsid protein VP1 to illustrate the diversity of new Microviridae viruses (Fig. 3). According to this analysis, virus 119 is closely related to unclassified Microviridae from moose (Alces alces), a close relative of the equine. Virus 348 indicates very small genetic change with bacteriophage Phi X 174, proposing subfamily Bullavirinae, based on the branch lengths.

**Figure 3:**
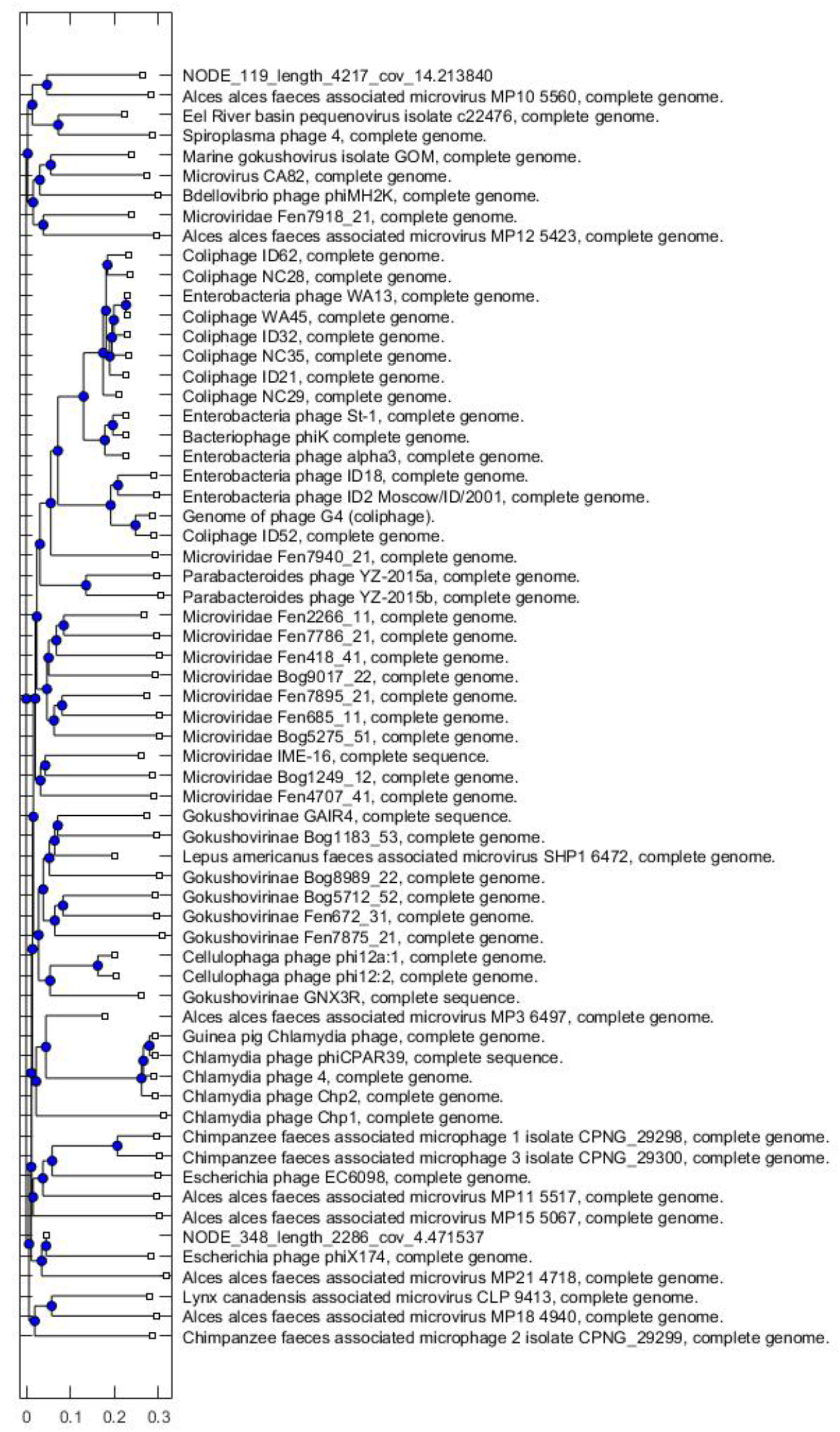
Maximum likelihood phylogenetic tree of full-length major capsid protein sequences present in the Microviridae genomes.

### Lifestyle and Host Prediction

Phages are broadly classified into two distinct lifestyles, lytic and temperate. With the prediction by DeePhage (Wu et al., 2021), the lifestyle for virus 119 is virulent (Supplementary 2), which can continuously complete five stages of attachment, genome insertion, biosynthesis, assembly and lysis in a short period of time to finish their proliferation. Virus 348 is temperate phage (Supplementary 2), which will integrate genome into the bacteria. When predicting the most likely bacterial host for the new phages, the most possible hosts for virus 119 is Candidatus Photodesmus blepharus (Supplementary 3). For virus 348, it is Alistipes putredinis (Supplementary 3).

### Comparative Analysis of Animals in qPCR and Sanger Sequencing

Results from sanger sequencing on both ends were matched with the sequence from assemble software for the equine, thus confirmed the assembly process of viral genomes from metagenomic reads used in this research and corroborated the obtained assemblies were not chimeric but represented as real viral genomes. When using the same primer sets to test other animals, novel viruses 119 and 348 are negative in canine and feline, ovine and porcine samples. However, virus 348 was identified positive in the bovine samples. Peak in melt curve overlap with the positive control. Besides, both forward and reverse primers can be found in sanger sequencing, which triple validated the accuracy of the sequence. Major capsid protein [Arizlama microvirus] was identified by blastx. The viral sequence shows 94.03% ident percentage alignment with the sequence in equine.

## Discussion

“One Health” concept means human health, animal health and our common environment are closely related. The rapid development of high-throughput DNA sequencing technology has demonstrated that many pathogens can be transferred between humans, animals, and the environment, which affects the research of entire microbial community (Trinh et al., 2018).

Compared with the human metaorganism, the complexity of the animal microorganism is often higher, because different animal species have unique microbes. In this research, we contributed in building the virome database by reporting 2 new complete virome sequences obtained from the equine uterine environment. In addition, we also justifiably suspect that there also exist within the bovine uterus different types of Microviridae, including one that appears to be closely related to the viral 348 genome that we discovered in equine. We also present indirect evidence suggesting that there are the same or similar Microviridae viruses in other animals who are genetically similar to the horse, like cattle and moose. Furthermore, virus 119 is very close to PhiX174, the very first organism to have the whole genome sequenced in 1977 (Sanger et al., 1977), and one of the main models in virology for genomic and capsid structure studies (Quaiser et al., 2015b). Nobel laureate researchers used it to prove covalently closed circularity (Fiers and Sinsheimer, 1962), while breaking through the age of synthetic biology (Goulian et al., 1967), identifying the role of enzymes in virus replication, complete oligonucleotides synthesize and particle assemble in vitro (Smith et al., 2003; Cherwa et al., 2011), and showing how the highly overlapping genome remain functional (Jaschke et al., 2012). With the high genetic similarity, virus 119 could offer another option to scientists under an era of rapid genetic evolution.

The ability of microorganisms to cooperate with each other makes them more stable and efficient in the ecosystem and empowers the microbiome with more comprehensive functions than a single microorganism. The host prediction provides us a new sight to think about the infectious disease treatment in the mare and cow uterus. One of the possible hosts, Alistipes putredinis, is a relatively newly identified bacteria isolated mostly from the GI system. Some teams have found it to be pathogenic in colorectal cancer and associated with depression, also being resistant to tetracycline and doxycycline (Parker et al., 2020). We could take advantage of the new phages to overcome the potential risk of antibiotic resistance. Therefore, future studies can focus on the role, these new viruses play in uterine environment.

Microorganisms are the main force of micro-environmental governance and remediation, and the basis of host function maintenance. Revealing the structure and function of the unknown virome and knowing the relationships between these new virome and different animal hosts are significant contents of microorganism learning.

